# Novel Potential *Cannabis* Pathogen: Discovery of Tobacco necrosis virus A in a Diseased Colombian *Cannabis sativa* Plant

**DOI:** 10.1101/2024.05.03.592441

**Authors:** Juliana Lopez-Jimenez, Herman D. Palacio-Torres, Juan F. Alzate

**Affiliations:** Centro Nacional de Secuenciación Genómica CNSG, Sede de Investigación Universitaria-SIU, Universidad de Antioquia, Medellín, Colombia; Fábrica de Plantas y Semillas de Antioquia S.A.S. – FASPLAN, El Carmen de Viboral, Antioquia, Colombia; Departamento de Microbiología y Parasitología, Facultad de Medicina, Universidad de Antioquia, Medellín, Colombia

**Author notes:** Correspondence: Juan F. Alzate, Departamento de Microbiologia y Parasitologia Facultad de Medicina, Centro Nacional de Secuenciación Genómica-CNSG Torre 1, S2-13, Sede de Investigación Universitaria-SIU Universidad de Antioquia, Carrera 53 # 61 - 30 Medellín. Colombia.

**Keywords:** RNA virus, TNVA, Tobacco Necrosis virus A, Transcriptome, *Cannabis sativa*, Hemp

## Abstract

Plant viral infections pose a significant threat to global crop productivity. Despite their profound impact on agriculture, plant viruses have been relatively understudied, primarily due to technological limitations associated with classical molecular methods. However, the advent of NGS RNA-seq analysis has revolutionized virus characterization in environmental settings, overcoming previous limitations and providing a powerful tool for studying plant viruses.

In an RNA-seq experiment conducted on a diseased Colombian *Cannabis sativa* hemp plant, we identified a linear single-stranded RNA (ssRNA) genome belonging to Tobacco Necrosis Virus A (TNVA), a common cause of necrotic lesions in plants such as tobacco and tulipa. The affected *Cannabis sativa* hemp plant exhibited severe symptoms, including alterations in pigmentation, leaf morphology such as chlorosis, necrotic tissue formation, and surface wear on the leaves. The complete genome sequence of the *Cannabis sativa* TNVA was 3,656 nucleotides long, containing five putative ORFs, and was classified in the family Tombusviridae, genus Alphanecrovirus, and belonging to the Necro-like clade based on RdRp protein phylogenetic analysis. Our analysis revealed a well-conserved RdRp protein among the Alphanecroviruses, with 89% of the amino acid residues in the peptide being entirely conserved. In contrast, the coat protein exhibited significantly higher variability, with only 49.3% of the residues being 100% conserved. Regarding the viral genome expression of *Cannabis sativa* TNVA, we observed that the virus was highly abundant in the leaves of the diseased plant, ranking among the topmost abundant transcripts, occupying the percentile position of 3.06%. Overall, our study generated the first reference genome of TNVA virus in the tropical region and reported the first case of this virus infecting a *Cannabis sativa* plant.

## INTRODUCTION

Tobacco necrosis virus A (TNVA) is an icosahedral virus that encapsidate one linear single-stranded RNA molecule ssRNA(+) and forms virions of 28 to 35 nm in diameter (Kotta-Loizou et al., 2019; Rubino & Martelli, 2008a). Its capsid is formed of 60 copies of a protein that forms trimers. Its genome has an average of 3.6 kb and 5 open reading frames (ORFs) that code for 5 different proteins (Garcia et al., 2022a).

According to International Committee on Taxonomy of Viruses (ICTV) at the taxonomic level, TNVA belongs to genus *Alphanecrovirus*, Tombusviridae family, Tolivirales order, Tolucaviricetes class, and Kitrinoviricota phylum. Recently, taxonomic adjustments have been made by changing the name of the specie tobacco necrosis virus A to *Alphanecrovirus nicotianae*. Until 2012, the genus Necrovirus was accepted, which was later divided into Betanecrovirus and Alphanecrovirus (C. M. Varanda et al., 2014a), from which seven different species arise. The family Tombusviridae has experienced adjustments and the addition of new genera in the recent past. The members of the genus Necrovirus have been reassigned as it is no longer in use (Monger & Jeffries, 2018a; Prasad et al., 2019)

Currently, *Tombusviridae* family is composed of three subfamilies: Procedovirinae, Regressovirinae and Calvusvirinae and eighteen genera such as *Alphacarmovirus, Alphanecrovirus, Aureusvirus, Avenavirus, Betacarmovirus, Betanecrovirus, Dianthovirus, Gallantivirus, Gammacarmovirus, Macanavirus, Machlomovirus, Panicovirus, Pelarspovirus, Umbravirus, Tombusvirus, Tralespervirus, Zeavirus* and *Luteovirus* (White, 2021; Zell et al., 2023). TNVA is taxonomically located within the Procedovirinae subfamily, known for its characteristic translational readthrough of a stop codon in the RdRp CDS sequence (White, 2021; Y. Morozov & G. Solovyev, 2020).

Four type species are members of the genus *Alphanecrovirus*: Tobacco necrosis virus A (TNVA), Olive latent virus 1 (OLV1), Olive mild mosaic virus (OMMV)(C. Varanda et al., 2018; C. M. Varanda et al., 2014b) and Potato necrosis virus (PoNV) (Hančinský et al., 2020a). An additional phylogenetic classification identifies a specific lineage known as the Necro-like clade. This lineage is derived from phylogenetic analyses based on the RdRp protein (Rubino & Martelli, 2021). Transmission of Necro-like viruses occurs through the soil and is thought to be facilitated by the presence of a vector such as the Chytridiomycetes fungus *Olpidium brassicae*, which spreads through their zoospores (Hančinský et al., 2020b; Rubino & Martelli, 2021; Teakle, 1960, 1962a).

Necro-like viruses are icosahedral virus composed of one linear positive sense single-stranded RNA genome, with an average size of 3.7 kb (Rubino & Martelli, 2021), that encode for five or six proteins, the 5’ end of the RNA is not capped, and the 3’ end does not have a poly (A) tail (Rochon, 1999; Shen & Miller, 2007). TNVA code for a pre-readthrough (pRT) RNA dependent RNA polymerase (RdRp), with the full length RdRp protein generated thanks to a readthrough of the stop codon UAG. They also encode the Coat protein (CP) (p30K), with an approximate size of 24–30 kDa protein, have been reported to interact with the zoospores of the fungal vector *Olpidium brassicae* (C. Varanda et al., 2011). Movement Protein 1 (MP1) (p8K) and Movement Protein (MP2) (p6K) are also encoded, these proteins allow their cell-cell movement. However, some viruses may have extra specific ORFs. TNVA contains five open reading frames (ORFs) and encodes five proteins, including pre-readthrough (pRT) RNA dependent RNA polymerase (RdRp) (ORF1), Movement Protein 1 (ORF2), Movement Protein 2 (ORF3), Coat protein (ORF4), and a hypothetical protein termed ORF5 (Rubino & Martelli, 2008b).

Necro-like viruses usually infects a very wide range of plants, including mono and dycotyledonous plants (Rubino & Martelli, 2021), including species of Amaranthaceae, Brassicaceae, Caprifolicaeae, Curcubitaceae, Fabaceae and Solanaceae families (Garcia et al., 2022b; Monger & Jeffries, 2018b; Verdin et al., 2018a; Zitikaitė & Staniulis, 2009). For research purposes, mechanical inoculation has been carried out on indicator plants such as *Nicotiana debneyi*, *Chenopodium amaranticolor* and *Chenopodium quinoa*. This process has resulted in the formation of localized necrotic lesions in inoculated leaves (Asjes, 1974), providing insight into its pathogenic behavior on crops (Monger & Jeffries, 2018a). TNVA natural infection seems to be concentrated in the roots of the plant, which is the place expected to be initially infected (King. et al., 2012a), however, systemic infection occurs in soybean *Glycine max* and *Nicotiana benthamiana* (Xi et al., 2008a; Y. Zhang et al., 2010). Natural infection by TNVA has been reported in China (Xi et al., 2008b), Hungary (Krizbai et al., 2010), New Zealand (Thomas & Fry, 1972), France (Verdin et al., 2018b) and Canada (Dias & Doane, 1968), however, some reports indicated a worldwide distribution. Tobacco necrosis virus A Chinese isolate (TNVA^C^) causes the necrosis and distortion of soybean, mulberry, potato and melon (Gao et al., 2021a; Xi et al., 2008b).

This study aims to achieve several goals: i) identify the possible etiology of plant deformation and the presence of chlorotic and necrotic lesions in a Colombian diseased *Cannabis* hemp plant, ii) gain a better understanding of TNVA infections in *Cannabis*, iii) generate the first TNVA genome sequence reference from *Cannabis*, and iv) raise awareness about the risk of TNVA infection in *Cannabis* crops.

## METHODS

### Sampling and Morphological characterization of plant lesions

The sampling process was carried out in the cultivation facilities of a company duly licensed by the Ministry of Justice, for the cultivation of *Cannabis sativa* plants with Psychoactive and Non-Psychoactive phytochemical profiles. The company is located in Cundinamarca department, in the center of Colombia, at an altitude of 2,566 meters above sea level. The cultivation of *Cannabis* plants takes place in greenhouses with plastic cover, division of areas, restricted access and process flows adjusted to a duly established system of good agricultural practices. This commercially significant variety, which exhibited signs and symptoms associated with alterations in its normal development, was selected and subsequently stabilized with trizol to avoid degradation of the genetic material.

### RNA-seq experiment on a Diseased Colombian Hemp Cultivar

Leaf samples were obtained from the diseased plant and immediately ground into powder using a mortar and liquid nitrogen. The powder was then suspended in Trizol reagent. RNA purification was performed following the manufacturer’s recommendations (TrizolTM Reagent, Invitrogen; RNA purification protocol). An acceptable RNA integrity was observed in the processed RNA sample with RIN value of 6.8. The RNA sequencing service was hired to Macrogen INC, a company based in Seoul, South Korea, using the Illumina NovaSeq6000 platform. rRNA-depleted libraries were prepared utilizing Illumina TruSeq kits, generating paired-end reads of 100 bp. The sequencing experiment produced 8,939,185,992 bases and 88,506,792 million paired end reads.

Read quality assessment and trimming were carried out using Cutadapt v3.5, with Q30 as the quality threshold (94,6) and a minimum length of 70 bases. The *de novo* transcriptome assembly was performed utilizing the Trinity 2.13.2 program (Grabherr et al., 2011). The contigs obtained from the assembly were employed to perform a search for TNVA genomes using the BLASTN tool version 2.12.0+ (Altschul et al., 1990; Z. Zhang et al., 2000). The database used was a custom database containing TNVA genome references downloaded from the GenBank. Subsequently, candidate sequences were chosen and manually verified to ensure the completeness of the viral genome sequence. Any incomplete sequences were ignored from further analysis.

### Annotation of the viral genome

A manual annotation of the complete sequence of the TNVA viral genome was made from the candidate assembled RNA-seq contigs. Manual annotation was done using the ARTEMIS tool version 18.0.2 (Rutherford et al., 2000) to validate its length, UTR sequences and the characteristics of the 5 CDS. Sequences classified as valid after manual verification were selected for further phylogenetic analysis. The viral genome was deposited on the NCBI database under the accession code PP584839.

### Phylogenetic analysis

We searched for nucleotide sequences of the complete genomes belonging to viruses of the *Alphanecrovirus* group within the NCBI public database. The dataset included seven genomes of the Olive latent virus 1 (OLV1) found infecting plant host *Nicotiana benthamiana* (NC_001721.1), *Solanum lycopersicum* (GU326337.2, OL472229.1, OL472228.1 and OR498642.1), *Olea europaea* (DQ083996.1) and *Nicotiana tabacum* (MK376952.1). Thirteen Olive mild mosaic virus (OMMV) found in *Valerianella locusta* (KY769774.1, KX906929.1), Tulipa (ON758350.1, MW241160.1, KU641010.1, KU641011.1), *Olea europaea* (NC_006939.1, HQ651834.1, HQ651832.1, , HQ651833.1), *Nicotiana benthamiana* (MW241164.1) and *Chenopodium quinoa* (MW241162.1). Beta vulgaris Alphanecrovirus 1 found in *Beta vulgaris* (MT227163.1), an *Alphanecrovirus* sp. in a River sediment metagenome (MZ218487.1), Potato necrosis virus (PoNV) found in *Solanum tuberosum* (NC_029900.1) and seven Tobacco necrosis virus A (TNVA) found in *Valerianella locusta* (KX906928.1), *Phaseolus vulgaris* (NC_001777.1), *Nicotiana benthamiana* (OP525286.1), *Glycine max (L.) Merr*. (AY546104.1) and *Chenopodium quinoa* (GQ221829.1 and MT675968.1) Table 1 Provides information about the *Alphanecrovirus* isolates in the dataset, which includes data as NCBI database access code, specie, isolate name, plant host, recollection country and date. Complete nucleotide sequences of 34 the viral genomes were subjected to a multiple alignment using the MAFFT software version 7.490 (Katoh & Standley, 2013). The final data set consisted of 32 *Alphanecrovirus* genomes and two betanecrovirus, Tobacco necrosis virus D (TNVD) genome sequences were used, found in *Nicotiana clevelandii* (NC_003487.1) and *Datura stramonium* (OL311682.1). The generation of phylogenetic trees was carried out with IQtree2 version 2.1.3 (Nguyen et al., 2015), including 5000 UFB pseudoreplicates. The substitution model that best suited the dataset was TIM2e+R4, according to model finder. TNVD nucleotide sequences were selected as outgroup. The graphic edition of the phylogenetic tree was carried out with the figtree program (FigTree version 1.3.1. Institute of Evolutionary Biology, University of Edinburgh, Edinburgh).

**Table 1.**
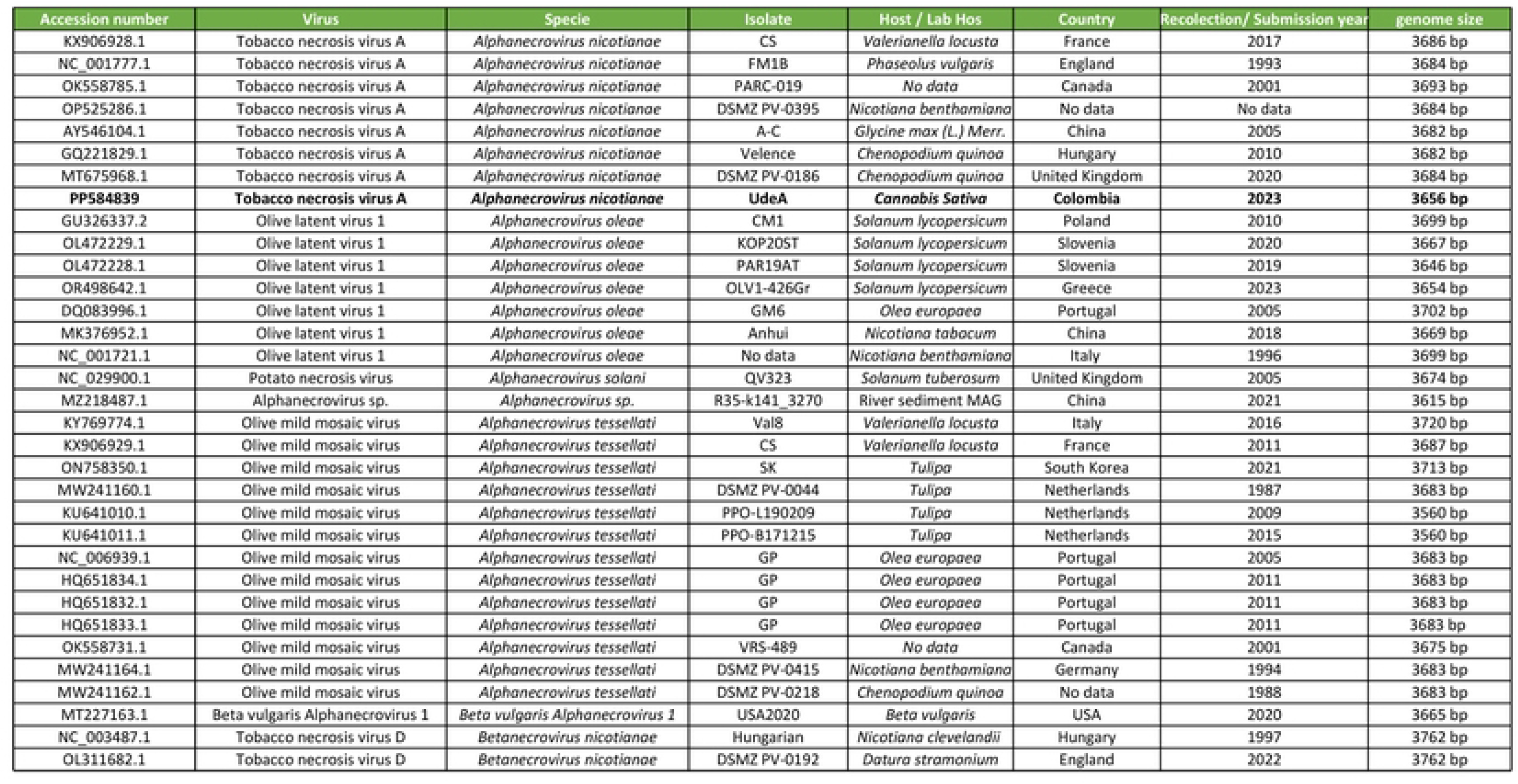
Information about the *Alphanecrovirus* in NCBI public database used in the phylogenetic analysis.

| Accession number | Virus | Specie | Isolate | Host / Lab Hos | Country | Recolection/ Submission year | genome size |
| --- | --- | --- | --- | --- | --- | --- | --- |
| KX906928.1 | Tobacco necrosis virus A | *Alphanecrovirus nicotianae* | CS | *Valerianella locusta* | France | 2017 | 3686 bp |
| NC_001777.1 | Tobacco necrosis virus A | *Alphanecrovirus nicotianae* | FM1B | *Phaseolus vulgaris* | England | 1993 | 3684 bp |
| OK558785.1 | Tobacco necrosis virus A | *Alphanecrovirus nicotianae* | PARC-019 | No data | Canada | 2001 | 3693 bp |
| OP525286.1 | Tobacco necrosis virus A | *Alphanecrovirus nicotianae* | DSMZ PV-0395 | *Nicotiana benthamiana* | No data | No data | 3684 bp |
| AY546104.1 | Tobacco necrosis virus A | *Alphanecrovirus nicotianae* | A-C | *Glycine max (L.) Merr.* | China | 2005 | 3682 bp |
| GQ221829.1 | Tobacco necrosis virus A | *Alphanecrovirus nicotianae* | Velence | *Chenopodium quinoa* | Hungary | 2010 | 3682 bp |
| MT675968.1 | Tobacco necrosis virus A | *Alphanecrovirus nicotianae* | DSMZ PV-0186 | *Chenopodium quinoa* | United Kingdom | 2020 | 3684 bp |
| **PP584839** | **Tobacco necrosis virus A** | ***Alphanecrovirus nicotianae*** | **UdeA** | ***Cannabis Sativa*** | **Colombia** | **2023** | **3656 bp** |
| GU326337.2 | Olive latent virus 1 | *Alphanecrovirus oleae* | CM1 | *Solanum lycopersicum* | Poland | 2010 | 3699 bp |
| OL472229.1 | Olive latent virus 1 | *Alphanecrovirus oleae* | KOP205T | *Solanum lycopersicum* | Slovenia | 2020 | 3667 bp |
| OL472228.1 | Olive latent virus 1 | *Alphanecrovirus oleae* | PAR19AT | *Solanum lycopersicum* | Slovenia | 2019 | 3646 bp |
| OR498642.1 | Olive latent virus 1 | *Alphanecrovirus oleae* | OLV1-426Gr | *Solanum lycopersicum* | Greece | 2023 | 3654 bp |
| DQ083996.1 | Olive latent virus 1 | *Alphanecrovirus oleae* | GM6 | *Olea europaea* | Portugal | 2005 | 3702 bp |
| MK376952.1 | Olive latent virus 1 | *Alphanecrovirus oleae* | Anhui | *Nicotiana tabacum* | China | 2018 | 3669 bp |
| NC_001721.1 | Olive latent virus 1 | *Alphanecrovirus oleae* | No data | *Nicotiana benthamiana* | Italy | 1996 | 3699 bp |
| NC_029900.1 | Potato necrosis virus | *Alphanecrovirus solani* | QV323 | *Solanum tuberosum* | United Kingdom | 2005 | 3674 bp |
| MZ218487.1 | Alphanecrovirus sp. | *Alphanecrovirus sp.* | R35-k141_3270 | River sediment MAG | China | 2021 | 3615 bp |
| KY769774.1 | Olive mild mosaic virus | *Alphanecrovirus tessellati* | Val8 | *Valerianella locusta* | Italy | 2016 | 3720 bp |
| KX906929.1 | Olive mild mosaic virus | *Alphanecrovirus tessellati* | CS | *Valerianella locusta* | France | 2011 | 3687 bp |
| ON758350.1 | Olive mild mosaic virus | *Alphanecrovirus tessellati* | SK | *Tulipa* | South Korea | 2021 | 3713 bp |
| MW241160.1 | Olive mild mosaic virus | *Alphanecrovirus tessellati* | DSMZ PV-0044 | *Tulipa* | Netherlands | 1987 | 3683 bp |
| KU641010.1 | Olive mild mosaic virus | *Alphanecrovirus tessellati* | PPO-L190209 | *Tulipa* | Netherlands | 2009 | 3560 bp |
| KU641011.1 | Olive mild mosaic virus | *Alphanecrovirus tessellati* | PPO-B171215 | *Tulipa* | Netherlands | 2015 | 3560 bp |
| NC_006939.1 | Olive mild mosaic virus | *Alphanecrovirus tessellati* | GP | *Olea europaea* | Portugal | 2005 | 3683 bp |
| HQ651834.1 | Olive mild mosaic virus | *Alphanecrovirus tessellati* | GP | *Olea europaea* | Portugal | 2011 | 3683 bp |
| HQ651832.1 | Olive mild mosaic virus | *Alphanecrovirus tessellati* | GP | *Olea europaea* | Portugal | 2011 | 3683 bp |
| HQ651833.1 | Olive mild mosaic virus | *Alphanecrovirus tessellati* | GP | *Olea europaea* | Portugal | 2011 | 3683 bp |
| OK558731.1 | Olive mild mosaic virus | *Alphanecrovirus tessellati* | VRS-489 | No data | Canada | 2001 | 3675 bp |
| MW241164.1 | Olive mild mosaic virus | *Alphanecrovirus tessellati* | DSMZ PV-0415 | *Nicotiana benthamiana* | Germany | 1994 | 3683 bp |
| MW241162.1 | Olive mild mosaic virus | *Alphanecrovirus tessellati* | DSMZ PV-0218 | *Chenopodium quinoa* | No data | 1988 | 3683 bp |
| MT227163.1 | Beta vulgaris Alphanecrovirus 1 | *Beta vulgaris Alphanecrovirus 1* | USA2020 | *Beta vulgaris* | USA | 2020 | 3665 bp |
| NC_003487.1 | Tobacco necrosis virus D | *Betanecrovirus nicotianae* | Hungarian | *Nicotiana clevelandii* | Hungary | 1997 | 3762 bp |
| OL311682.1 | Tobacco necrosis virus D | *Betanecrovirus nicotianae* | DSMZ PV-0192 | *Datura stramonium* | England | 2022 | 3762 bp |

### Cannabis sativa leaves RNA-seq raw data from the SRA-NCBI database

*Cannabis sativa* leaf RNA-seq data from the SRA-NCBI database was acquired in November 2021 and subsequently assembled using Trinity, following the methodology outlined previously. A total of 107 transcriptomes from *Cannabis sativa* leaves were examined for the presence of the TNVA virus, mirroring the approach described earlier for the Colombian hemp plant. Remarkably, only one transcriptome, originating from a *Cannabis* plant in China (SRA accession code: SRR2961016), yielded a significant hit against the TNVA virus. Subsequent manual inspection and annotation of the viral genome contig ensured its quality, and classification was carried out using the same phylogenetic methodology.

### Tobacco Necrosis Virus A genome expression analysis

Relative abundance analysis of the TNVA viral genome was conducted using the Kallisto package v0.44.0 (Bray et al., 2016), employing the TPM (Transcripts Per Million) normalized metric. In this analysis, the longest contig of each assembled transcript variant, including its respective viral genome contigs, was chosen as the reference sequence. Kallisto aligned and counted the transcriptomic clean reads against these references, subsequently calculating the TPM values. The resulting dataset was imported into the R environment for further manipulation and graphical processing using the ggplot2 library.

## Data availability

The Colombian TNVA genome sequence generated in this work was deposited at the NCBI GenBank database under the accession code: PP584839.

## Funding

Fundación Universidad de Antioquia.

Universidad de Antioquia, Centro Nacional de Secuenciación Genómica-CNSG.

## RESULTS

### Description of the Diseased *Cannabis sativa* hemp plant

There are several alterations that can be observed in the leaf structure of the plant. One of the main aspects to highlight is the presence of eroded segments, both on the upper and lower sides of the leaves, characterized by a light brown color. These effects on the leaf surface, possibly generated by the presence of scraping insects, increase the level of vulnerability of the plant to potential infections, due to the exposure of the vascular system. On the other hand, changes can be observed in the natural pigmentation of the plant, going from dark green to yellow, a phenomenon known as chlorosis. In the particular case of the plant evaluated, this chlorosis is distributed heterogeneously in some leaves, however, the presence of a completely chlorotic segment stands out, in the upper part of the central leaflet of one of the leaves. It should be noted that this major chlorosis is delimited by a cord of necrotic damage that extends perpendicular to the veins of the leaflet, evident by its dark brown color and the alteration of symmetry in the tissue. These signs and symptoms somehow show physiological alterations that affect the correct development of the plant (Figure 1).

**Figure 1.**
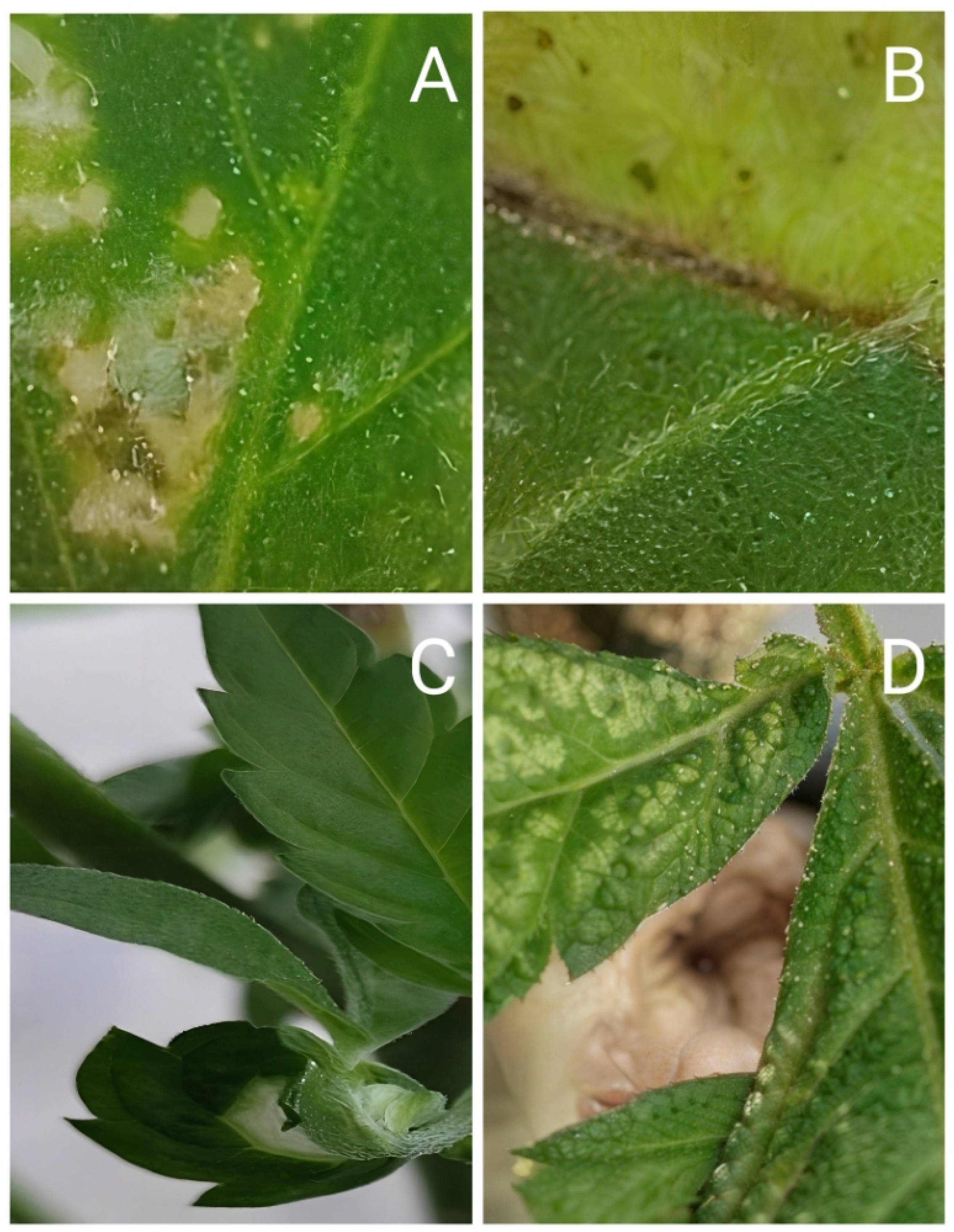
Morphological characteristics of cannabis plant infected with TNVA. Different types of alterations at the level of leaf pigmentation and leaf morphology can be seen. Figure 1A highlights the presence of small eroded segments, which turn brown, typical of an oxidative deterioration process. Figure 1B shows chlorosis in the upper part of the central leaflet of the leaf. This alteration is marked by the formation of a necrotic tissue that separates the pigmented leaf segment from the chlorotic segment. Figure 1C indicates the abnormal development of a leaf marked by the twisting and the absence of half a leaf segment. Figure 1D shows deterioration and wear on the leaf surface, usually attributable to the presence of small insects.

**Discovery of Tobacco Necrosis Virus A in Diseased Colombian *Cannabis sativa* Hemp Plants** In pursuit of identifying the etiological agent responsible for the observed disease in the aforementioned plants, we conducted a detailed transcriptome analysis to search for viral agents that might be replicating within the leaves displaying lesions in the diseased Colombian *Cannabis sativa* plant. To accomplish this, total RNA was extracted from the diseased leaves, and an rRNA-depleted RNA-seq library was prepared and sequenced. After filtering for read quality and *de novo* assembly, contigs containing nucleotide sequences of known plant viruses were examined. In the diseased Colombian *Cannabis sativa* plant, we identified one contig carrying the entire viral genome of the *Cannabis sativa* Mitovirus CasaMV1 lineage 2 (Lopez-Jimenez et al., 2023).

Additionally, we detected a contig spanning 3,656 bases that, upon initial evaluation, encompassed all the five coding sequences (CDS) of the TNVA virus, with an E-value of 0 and a nucleotide identity percentage of 96.7% to the TNVA genome reference (GenBank Accession KX906928). The TNVA genomic contig showed an average sequencing depth of 126.7X. Manual annotation of this transcriptomic contig confirmed the presence of the complete five expected CDSs of the TNVA viral genome: ORF1, ORF1-RT (RNA-dependent RNA polymerase - RdRp), ORF2 (Movement protein 1), ORF3 (Movement protein 2), ORF4 (Coat protein), and ORF 5 (Hypothetical protein). This annotation process involved validating the presence of the start ATG codon, confirming the length of the CDS, and ensuring the presence of the respective stop codons (Figure 2).

**Figure 2.**
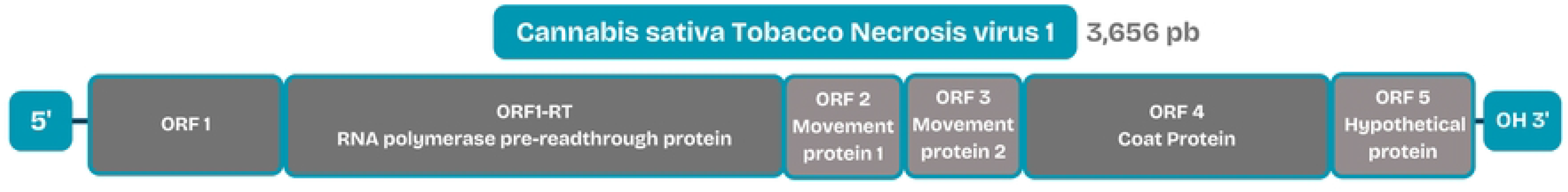
Genome structure of the *Cannabis sativa* Tobacco necrosis virus A (TNVA) and its coding sequences. This schematic representation illustrates the genome structure of the Colombian *Cannabis sativa* isolate of the Tobacco necrosis virus A (TNVA), highlighting the positions of its coding sequences (CDSs). The viral genome consists of a linear single-stranded RNA molecule, spanning 3,656 bases. The coding sequences (CDSs) are depicted as gray boxes along the genome, each representing a distinct gene responsible for encoding specific viral proteins. The BOX lengths are proportional to the CDS length.

***Cannabis sativa* TNVA taxonomic confirmation, *Alphanecrovirus* evolution and host range** With the aim of analyzing the evolutionary history of the reported genomes of the *Alphanecroviruses* and also to confidently confirm the taxonomic assignment of the putative *Cannabis sativa* TNVA, a maximum likelihood phylogenetic tree of the genus *Alphanecrovirus* was computed. To achieve this, a nucleotide alignment matrix encompassing the complete genomes of the reported *Alphanecroviruses* isolated from plants, including the Colombian *Cannabis* TNVA isolate, was generated and uploaded into IQTREE2 to construct the tree. The resulting tree reconstructs the genus *Alphanecrovirus* with robust support (99%). Additionally, the tree topology depicts OLV1 and TNVA as sister clades, both forming well-supported monophyletic clades for each virus taxon. Remarkably, within the TNVA clade, the well-supported position of the Colombian *Cannabis sativa* TNVA confirms its taxonomic assignment as Tobacco Necrosis Virus A (Figure 3).

**Figure 3.**
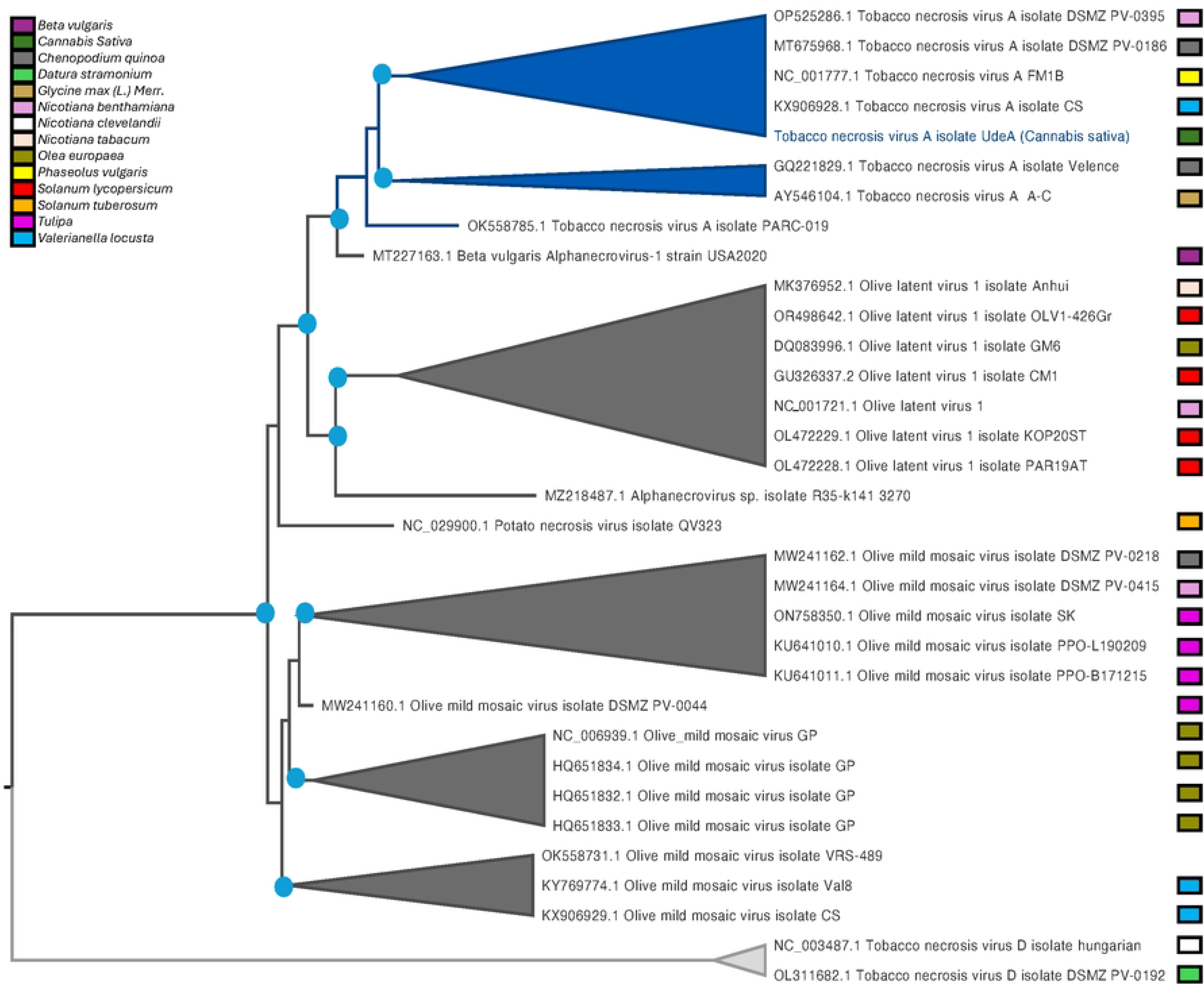
Phylogenetic Analysis of Alphanecroviruses TNVA, OMMV, and OLV1 Isolates from Plants. Viral genome maximum-likelyhood phylogenetic tree portraying the evolutionary relationships among different isolates of the Alphanecroviruses Tobacco necrosis virus A (TNVA), Olive mild mosaic virus (OMMV), and Olive latent virus (OLV) isolated from plants. Blue circles denote UFB bootstrap values above 95% (>95) for its respective branch. The selected outgroup was TNVD.

We sought to compare the host plants from which the Alphanecroviruses were isolated, aiming to better understand the virus-host plant spectrum. As illustrated in the phylogenetic tree color-coded, Olive mild mosaic virus exhibits a broad spectrum of host plants, spanning five orders: *Nicotiana* (Solanales), *Chenopodium* (Caryophyllales), *Olea* (Lamiales), *Valerianella* (Dipsacales), and *Tulipa* (Liliales) (Figure 3). In contrast, OLV-1 displays a narrower spectrum, primarily isolated from *Solanaceae*, with one exception isolated from Lamiales (*Olea*). The TNVA virus demonstrates a wide host plant spectrum, isolated from plants belonging to the orders Fabales, Caryophyllales, Urticales, Dipsacales, and Solanales (Table 1) (Figure 3).

We also aimed to explore the geographical origin of the reference *Alphanecrovirus* genomes deposited in GenBank. This analysis indicates that the majority of these viral genome references originate from studies conducted in developed northern hemisphere temperate countries, with genome references reported from North America, Western Europe, and Asia. The Colombian *Cannabis sativa* TNVA isolate represents the first report from a tropical region and also the first report in a *Cannabis* plant (Figure 4).

**Figure 4.**
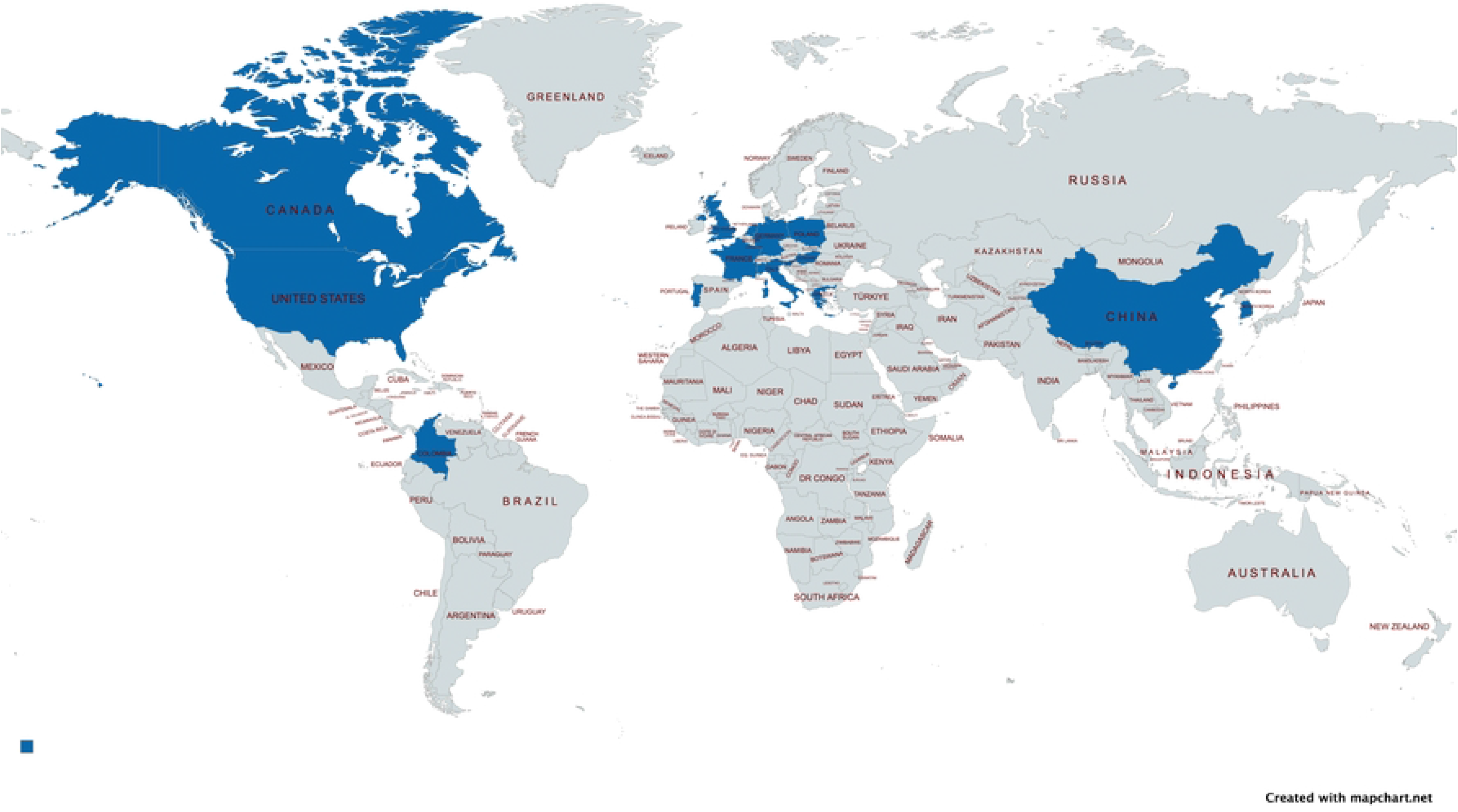
Countries Reporting Genome Sequences of the genus *Alphanecrovirus*. This world map illustrates countries where *Alphanecrovirus* genome sequences isolated from infected plants have been reported. Countries with viral genome references documented in GenBank are highlighted in blue. Remarkably, Colombia is the first tropical country to report naturally infected plants, and it also marks the first report of natural infection in a *Cannabis* hemp plant.

### RNA dependent RNA polymerase and viral Coat Protein conservation in TNVA

RNA dependent RNA polymerase and Coat proteins are extremely important for viral replication and transmission, respectively. To assess the conservation of their respective coding sequences (CDSs) and peptides, we aligned and analyzed them. The RdRp CDS spanned 2,172 nucleotides in all reference genomes, with no gaps present in any sequence. The CDS was highly conserved, with 77.7% of nucleotide residues being 100% conserved, while 1.9% of the nucleotides showed conservation percentages equal to or lower than 50% (Figure 5, panel A).

**Figure 5.**
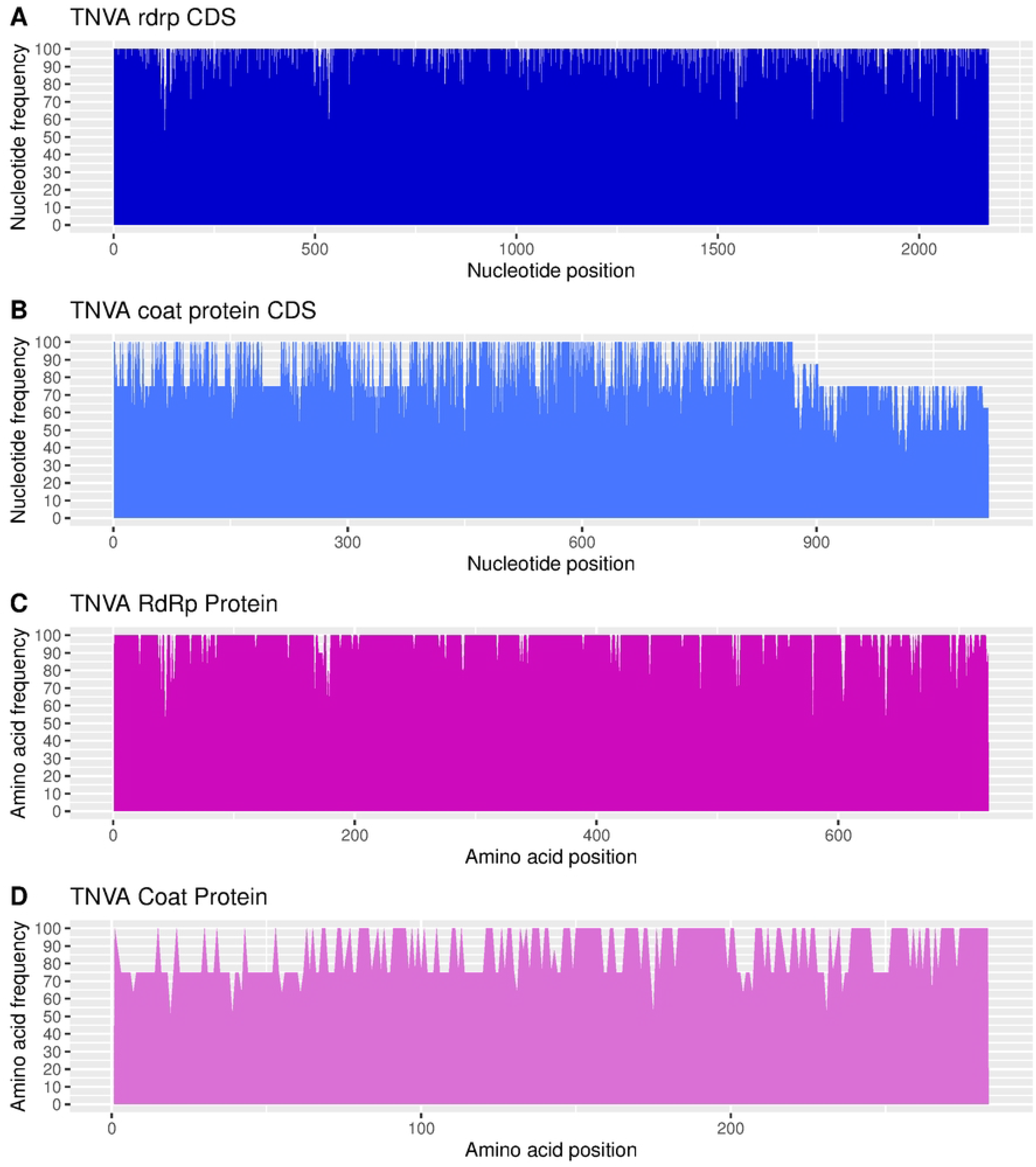
Conservation Patterns of Genes and Proteins in Tobacco Necrosis Virus A (TNVA) This figure illustrates the conservation patterns of genes and proteins, specifically the RdRp polymerase and the coat protein, in Tobacco Necrosis Virus A (TNVA). Panel (A) and Panel (B) present area plots showing the frequency distribution of the most conserved nucleotide across the entire gene of the RdRp polymerase and the coat protein coding sequences (CDS) of TNVA, respectively. The x-axis represents the position of the nucleotide, while the y-axis denotes the frequency of the most common nucleotide at each position. In Panels (C) and (D), the area plots depict the conservation patterns across the RdRp polymerase (C) and the coat protein (D) of TNVA at the amino acid level. Here, the x-axis indicates the position of the amino acid residues, while the y-axis displays the frequency percentage of the most prevalent amino acid.

For the coat protein CDS, a less conserved profile was observed, with a typical length of 828 bases (276 amino acids) and the presence of gaps in the alignment and significant variations, especially at the 5’ regions of the CDSs. Approximately 1.8% of the of nucleotide positions showed low conservation (≤ 50%), while 50.5% were 100% conserved (Figure 5, panel B).

The conservation of the respective viral proteins was notably higher, particularly in the RdRp protein, where 89% of the amino acid residues in the peptide were entirely conserved.

Additionally, all predicted peptides displayed the same amino acid length, 724 residues. Only three amino acid positions showed conservation values equal to or lower than 50% (Figure 5, panel C). In contrast, the coat protein, typically comprising 276 amino acids, exhibited significantly higher variability, with only 49.3% of the residues being 100% conserved (Figure 5, panel D).

### Viral genome expression of the Cannabis TNVA

Finally, we wanted to test how abundant was the *Cannabis sativa* TNVA virus within the transcriptome of the *Cannabis* hemp diseased plant. To do so, we quatified the relative abundance of each transcriptomic contig using the Transcripts Per Million (TPM) units and plotted its length in bases versus its relative abundance in TPMs. For visualization enhancement, the contig with the viral genome of the *Cannabis sativa* TNVA virus is depicted as a red triangle (Figure 6, panel A). As depected in the figure, the *Cannabis sativa* TNVA is highly abundant, supported by 2,255 high quality reads, hence highly expressed, being as one of the top most abundant transcripts in the leaves of the diseased hemp plant occupying the percentile position 3.06%.

**Figure 6.**
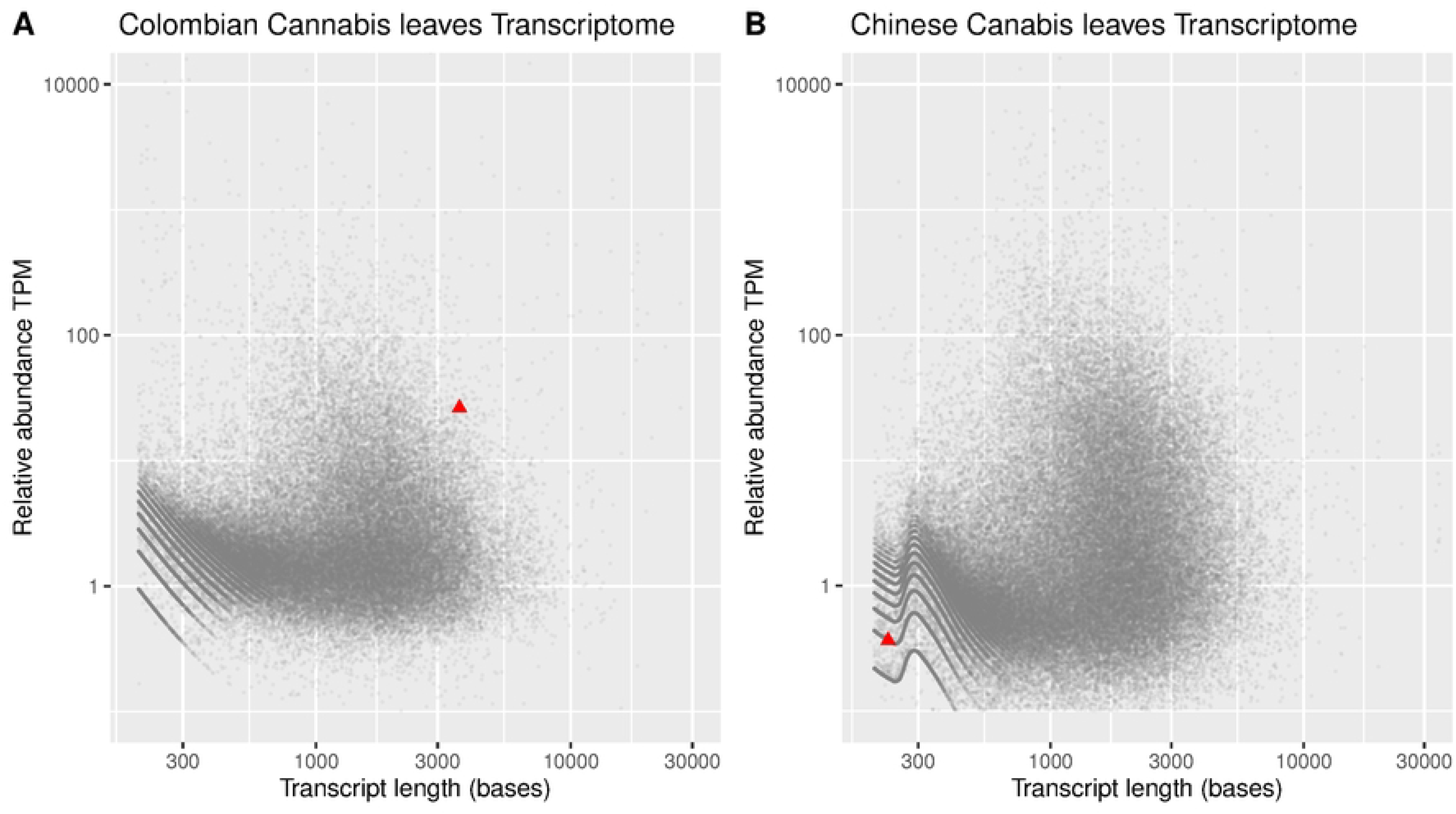
Transcriptome Expression Levels of Tobacco Necrosis Virus A - TNVA in *Cannabis sativa* leaves. This figure presents scatter plots illustrating the *de novo* assembled contigs of the transcriptome of *Cannabis sativa* leaves. Each contig is represented as a gray dot, with the y-axis indicating the relative abundance in TPM (Transcripts Per Million) units and the x-axis denoting the transcript length in bases. Both axes utilize a logarithmic scale for improved visualization. Red triangles indicate contigs corresponding to the TNVA viral RNA genome. Panel A depicts the leaf transcriptome of a Colombian *Cannabis sativa* plant infected with TNVA, while Panel B showcases the leaf transcriptome of an infected Chinese hemp plant found in the SRA database (SRR2961016).

To complement these results, we utilized a local custom database comprising 107 transcriptomes of *Cannabis sativa* leaves (Lopez-Jimenez et al., 2023) to search for other plants infected with the TNVA virus. Remarkably, in one transcriptome from leaves of a hemp plant cultivated in China (SRA accession code: SRR2961016), we identified a partial sequence of the TNVA viral genome, consisting of a contig with 228 bases and exhibiting 95.5% nucleotide identity with the Colombian *Cannabis* TNVA. Phylogenetic analysis confirmed the taxonomic assignment to the TNVA clade with robust support (see Supplementary Figure 1). Analysis of relative abundance within its respective transcriptome revealed that in the leaves of this hemp plant, the TNVA virus was expressed lightly, positioned at 80.32% in the percentile rank (Figure 6, panel B).

## DISCUSSION

Viral infections in plants represent a significant threat to the development and productivity of crops, resulting in substantial losses worldwide (Rubio et al., 2020; Tatineni & Hein, 2023). Despite their profound impact on global agriculture, plant viruses remain poorly studied, especially in developing nations. The emergence of new viral diseases results from global trade, climate fluctuations, and viruses’ capacity for quick evolution (Rubio et al., 2020). Viruses present unique challenges for detection, primarily due to their minute size, at the nanometer scale, and the inability to culture them axenically due to their highly specialized parasitic nature. Classical detection methods are often limited by technological constraints, underscoring the need for advanced technologies such as Next-Generation Sequencing (NGS) to monitor plant viral infections effectively. NGS technologies offer unparalleled capabilities for surveillance, particularly in dynamic viral landscapes where novel viruses or variants emerge regularly through evolution (Mehetre et al., 2021; Tatineni & Hein, 2023). NGS technologies does not require any previous knowledge of viral sequences, allowing the detection of viruses that affect a plant, both known and unknown (Rubio et al., 2020).

Several factors contribute to the rapidly deteriorating global situation of plant virus diseases, like the expansion of international trade by companies introducing damaging virus diseases to regions where they were previously absent. This is driven by trade globalization and more efficient transport methods. Additionally, the movement of crop plants away to distant regions results in the emergence of new virus diseases at an accelerating rate. Plant virus disease are becoming progressively harder to manage, mainly for climate instability and global warming. An additional risk is the seed trade, where it is likely to introduce seed-borne viruses (Jones, 2021).

The transmission of plant viruses through water represents a major concern for sustainable food and agricultural systems. Many plant pathogenic viruses, particularly those belonging to the Tombusviridae family, have been recovered from environmental waters. The ability of these viruses to be transmitted through the environment depends on how long they can survive in both terrestrial and aquatic environments. Over recent decades, several epidemics caused by new viruses that spilled over from reservoir species or new variants of classic viruses have had detrimental effects on global agricultural production (Betancourt, 2023).

The family Tombusviridae includes a diverse array of soil and water-borne viruses that infect plants and are distinguished by their icosahedral virions measuring approximately 30 nm in diameter and a compacted single-stranded positive-strand RNA genomes (Scheets et al., 2015). The main criteria for the taxonomic classification of Tombusviridae family is the structural relationship between their viral supergroup-II RdRps, which exhibit high sequence similarity (Castaño & Hernández, 2005; Nicholson et al., 2012). The significant level of diversity among species of Movement Proteins and silencing suppressor encoded, have also been used as a guide for taxonomic classification. Phylogenetic analyses based on RdRps provide insights into the evolutionary history of these viruses, revealing several distinct lineages such as Tombusvirus-like, Necrovirus-like, and Carmovirus-like. Within these lineages, the genera Alphanecrovirus, to which TNVA belongs, is situated within the Necro-like lineage. Additionally, genera such as Tombusvirus, Aureusvirus, Zeavirus, and Betanecrovirus are found within the Tombusvirus-like lineage (King. et al., 2012a).

Tombusviridae viruses infect monocotyledonous and dicotyledonous plants (Agrios, 2005; King. et al., 2012b). Experimental inoculations usually cause necrotic lesions on the inoculated leaves, but rarely result in systemic infection of the plant (Garcia et al., 2022b). TNVA are widely distributed and their virions are readily transmitted by mechanical inoculation. These viruses persist in the environment for long periods. Some research suggests that *Olpidium brassicae*, a chytrid fungus, might play a role in naturally transmitting TNVA (Teakle, 1960, 1962b), but the exact mechanisms and significance of this transmission route are still being investigated. Olive latent virus-1 (OLV-1), another member of the Tombusviridae family, seems to be transmitted through the soil without the apparent intervention of a vector (King. et al., 2012b) The first report of Tobacco necrosis virus (TNVD) in Colombia occurred recently in 2023, specifically pinpointing TNVD as the identified virus in potato seed-tubers originating from the northern Antioquia region, nestled within the Andean landscape of Colombia. This study utilized NGS RNA-seq technology in *Solanum phureja* and facilitated the retrieval of the partial sequence of the TNVD genome (García et al., 2023). TNVD is classified within the genus Betanecrovirus (Rochon, 1999; C. M. Varanda et al., 2014b; Verdin et al., 2018a), standing as a sister clade to the Alphanecrovirus. The Betanecrovirus genus comprises two more species: Beet black scorch virus (BBSV) and Leek white stripe virus (LWSV) (Chkuaseli et al., 2015).

In our study, we were able to recover a nearly full-length TNVA viral genome from a *Cannabis* hemp plant cultivated as an industrial crop in the Andean region of Colombia. The transcriptomic contig assembled in our investigation encompassed all five reported CDS sequences and was supported with a confident average sequencing depth of 126X. This represents not only the inaugural report of this virus infecting the genus *Cannabis* but also marks the first documentation of a reference viral genome for TNVA in the tropical region of the planet.

Our study also aimed to elucidate the host plant spectrum of Alphanecroviruses to gain a deeper understanding of their ecological and agricultural significance. Our phylogenetic analysis revealed distinct patterns in the host range among different Alphanecroviruses. Specifically, Olive mild mosaic virus displayed a wide spectrum of host plants, spanning across five taxonomic orders including Solanales, Caryophyllales, Lamiales, Dipsacales, and Liliales. This broad host range suggests a high degree of adaptability and versatility in its interactions with various plant species.

In contrast, OLV-1 exhibited a narrower host spectrum, primarily infecting plants within the Solanaceae family, with a single exception observed in the Lamiales order (*Olea*). This restricted host range may reflect specific evolutionary adaptations or ecological constraints governing the virus-plant interactions of OLV-1. Interestingly, the TNVA virus demonstrated a remarkably wide host plant spectrum, infecting plants from diverse taxonomic orders including Fabales, Caryophyllales, Urticales, Dipsacales, and Solanales. This broad host range underscores the potential agricultural impact and ecological implications of TNVA infections across a wide range of plant species. Overall, our findings contribute to a comprehensive understanding of the host range dynamics of Alphanecroviruses and highlight the importance of considering host plant diversity in virus ecology and management strategies.

Our analysis of the RdRp and the virus coat protein highlights the critical role of the former in maintaining viral genome integrity, as evidenced by its lower variation, even at the nucleotide level. This highlights the essential nature of the RdRp protein and suggests strong evolutionary constraints on this region. In contrast, the coat protein CDS exhibited a less conserved profile, with notable variability observed among the tested reference viral genomes. This greater variability in the coat protein may indicate potential adaptability to diverse environmental conditions or hosts.

The abundance of the TNVA virus within the transcriptome of diseased *Cannabis sativa* hemp plants was a key focus of our investigation, particularly considering its potential association with the observed plant lesions. Our analysis of relative abundance underscores the significant prevalence of the TNVA genome within the infected leaf transcriptome, indicating robust transcriptional activity and potentially inducing cellular stress in the infected plant cells. While we refrain from definitively establishing a direct causal link between TNVA and the observed plant disease, the exclusive presence of this virus within the affected leaves prompts further inquiry. Future studies are warranted to elucidate whether TNVA indeed plays a central role in the observed disease manifestation in *Cannabis* plants.

The search for TNVA presence in *Cannabis* plants across different latitudes led us to detect an infected hemp plant cultivated in China. Despite the genetic similarity observed between TNVA viruses isolated from disparate regions such as Colombia and China, the expression level within the transcriptome of the Chinese hemp plant was notably lower. This significant difference in expression levels may suggest variations in viral replication dynamics or host-virus interactions. However, it’s crucial to note that we lack information regarding the health status of the Chinese plant, preventing us from extending our conclusions to whether similar disease symptoms were observed in that plant.

In previous studies, TNVA has been observed to infect soybeans systemically under natural conditions, leading to necrosis and distortion of soybean plants as early as 1982 (Xi et al., 2008a). When TNVA is introduced through mechanical inoculation, the infection triggers a localized necrotic response, primarily attributed to the hypersensitive reaction (HR) initiated by the virus coat protein. Additionally, systemic chlorotic symptoms manifest as a result of the infection.

These distinct lesions typically emerge within 3–4 days post-infection and offer a quantifiable indicator of viral activity. Consequently, this pathosystem has been widely utilized to evaluate both the extent and underlying mechanisms of induced plant resistance against viral infections (Gao et al., 2021b).

In conclusion, our study underscores the global prevalence of the TNVA virus and its ability to infect *Cannabis sativa* plants. Our findings provide preliminary evidence supporting the potential pathogenicity of TNVA in *Cannabis*. However, further research is imperative to definitively establish its pathogenic behavior in *Cannabis sativa* crops. Therefore, we recommend including TNVA in the list of potential pathogens for *Cannabis* sativa and advocate for continuous monitoring by breeders to detect its presence in diseased plants. Building a comprehensive body of knowledge over time will facilitate a more accurate assessment of TNVA’s impact on plant health and agricultural productivity.

---

**Supplementary Figure 1 Phylogenetic Analysis of Alphanecroviruses including TNVA isolated from Chinese Cannabis Plant.**

Viral genome maximum-likelyhood phylogenetic tree portraying the evolutionary relationships among different isolates of TNVA, including two *Cannabis sativa* TNVA.

